# Subcellular micropatterning for visual immunoprecipitation reveals differences in cytosolic protein complexes downstream the EGFR

**DOI:** 10.1101/2021.05.25.445547

**Authors:** Roland Hager, Ulrike Müller, Nicole Ollinger, Julian Weghuber, Peter Lanzerstorfer

## Abstract

Analysis of protein-protein interactions in living cells by protein micropatterning is currently limited to the spatial arrangement of transmembrane proteins and their corresponding downstream molecules. Here we present a robust method for visual immunoprecipitation of cytosolic protein complexes by use of an artificial transmembrane bait construct in combination with micropatterned antibody arrays on cyclic olefin polymer (COP) substrates. The method was used to characterize Grb2-mediated signalling pathways downstream the epidermal growth factor receptor (EGFR). Ternary protein complexes (Shc1:Grb2:SOS1 and Grb2:Gab1:PI3K) were identified and we found that EGFR downstream signalling is based on constitutively bound (Grb2:SOS1 and Grb2:Gab1) as well as on agonist-dependent protein associations with transient interaction properties (Grb2:Shc1 and Grb2:PI3K). Spatiotemporal analysis further revealed significant differences in stability and exchange kinetics of protein interactions. Furthermore, we could show that this approach is well suited to study the efficacy and specificity of SH2 and SH3 protein domain inhibitors in a live cell context. Altogether, this method represents a significant enhancement of quantitative subcellular micropatterning approaches as an alternative to standard biochemical analyses.

## Introduction

Protein micropatterning has become an important tool for fundamental research in cell biology. Micropatterned substrates were mainly engineered for the investigation of the influence of the extracellular environment on cell morphology, cell differentiation, cytoskeleton rearrangement, cell migration and organelle positioning (Théry, 2010; Albert and Schwarz, 2016). Within this regard, soft lithography via microcontact printing (μCP) is one of the most convenient and widely used methods for patterning proteins on a micron- and even nanometer-scale (Perl et al., 2009; Kaufmann and Ravoo, 2010; Lindner et al., 2018).

Recently, we and others have developed protein micro- and nanopatterning approaches on solid substrates for the quantitative investigation of protein-protein interactions (PPIs) in the live cell (Lanzerstorfer et al., 2014; Löchte et al., 2014; Lanzerstorfer et al., 2015; Wakefield et al., 2017; Dirscherl et al., 2017; Schütz et al., 2017; Lindner et al., 2018; Dirscherl et al., 2018; Motsch et al., 2019; Lanzerstorfer et al., 2020). The fundamental strategy of these approaches is the spatial rearrangement of a cell surface protein (‘bait’, e.g. receptor) in the shape of the printed patterns within the plasma membrane (e.g. by use of antibodies or ligands) and the monitoring of the lateral distribution of a putative interaction partner (‘prey’, e.g. cytosolic adapter protein). This enables the investigation of PPIs in a native environment and membrane composition, and importantly, allows for straight-forward characterization of PPIs in the living cell. However, these methods are mostly limited to protein complexes formed between a membrane-anchored bait and an intracellular prey.

As cytosolic protein complexes play a key role in the precise regulation of cellular signalling events, they have become putative new selective drug targets (Lee et al., 2019). Hence, there is an increasing interest in the design and development of robust experimental approaches beyond standard biochemical methods (e.g. such as co-immunoprecipitation, pull-down experiments and yeast two-hybrid screens) that enable in-depth characterization of protein complexes inside cells. Information on interaction properties such as binding affinities, lifetimes, stability, and dynamics of protein complex formation is of particular importance, as these parameters are critical for the regulation of cellular systems (Perkins et al., 2010).

Within this regard, an in situ single-cell pull-down approach on micropatterned functionalized surface architecture in combination with single-molecule fluorescence imaging was developed to measure protein complex stoichiometry and dynamics (Wedeking et al., 2015). Recently, a microfluidic device for in situ co-immunoprecipitation of target proteins to detect PPIs in individual cancer cells with high fidelity was reported (Ryu et al., 2019). Furthermore, a singlemolecule pull-down (SiMPull) assay was described which enables direct visualization of individual cellular protein complexes by single-molecule fluorescence microscopy (Jain et al., 2011). However, those sophisticated approaches have in common that cells cultured on these supports need to be lysed by detergents prior to PPI analysis and therefore do not allow for live cell measurements. In order to analyze PPIs in the cytosol of living cells, Gandor et al. (2013) reported on a strategy using artificial receptor constructs (termed bait-PARCs) that transfer a micrometer-scale antibody surface pattern into an ordered array of cytosolic bait proteins in the plasma membrane. Most recently, a similar approach for real-time quantification of cytosolic PPIs using cell-based molography as a biosensor was developed (Incaviglia et al., 2020). In addition, subcellular micropatterning of artificial transmembrane receptors was proofed by using fibrinogen anchors (Watson et al., 2021).

Based on the most recent developments, which also demonstrate the importance and future applicability of micropatterned interfaces for intracellular PPI analysis, we here report on a robust platform for visual immunoprecipitation of cytosolic PPIs. The approach is based on subcellular micropatterning of bait-presenting artificial transmembrane constructs in the cytoplasm of living cells. We introduced cyclic olefin polymer (COP) as a cost-saving and flexible alternative to glass coverslips for large-area microcontact printing and realized a 384-well plate-based platform with modular protein micropatterns which enabled an increased experimental throughput. Furthermore, we redesigned and optimized bait-presenting artificial receptors (herein referred to as bait-PARs) for enhanced prey corecruitment. In order to demonstrate the applicability of the method, we investigated cytosolic protein complexes downstream the epidermal growth factor receptor (EGFR). The EGFR has become one of the most extensively studied cell surface receptors and a major oncogenic drug target, as aberrant receptor activation and intracellular signal transduction is associated with a variety of cancers, thus making its key players in downstream signalling to the perfect proof of concept target for our study (Tomas et al., 2014). We could unequivocally show that EGFR downstream signalling is based on Grb2-mediated ternary protein complexes exhibiting different interaction regimes (constitutively bound vs. agonist-dependent). Additionally, we identified significant differences in protein complex formation kinetics and stability of detected assemblies, which might account to the dynamic regulation of normal and aberrant EGFR signalling. Furthermore, we characterized the efficacy and specificity of therapeutic Src Homology (SH) domain inhibitors with high sensitivity.

Altogether, we could demonstrate that our technology allows for the control of the subcellular localization of cytosolic adapter proteins, hence enabling the spatiotemporal investigation of receptor-mediated intracellular PPIs within a defined signalling cascade. With the introduction of this robust and flexible assay, we introduce an add-on to standard biochemical PPI analysis, which might facilitate protein micropatterning for cell biological investigations in the future.

## Results and Discussion

### Fabrication of micropatterned COP foils using large-area microcontact printing (μCP)

We have recently introduced large-area patterned glass substrates with modular protein micropatterns that enable the systematic investigation of specific and nonspecific effects in the analysis of PPIs in adherent cells (Lanzerstorfer et al., 2020). However, functionalized glass substrates possess major drawbacks such as increased specific costs and high fragility, especially when used in combination with sensitive fluorescence spectroscopy approaches such as TIRF microscopy, as they require a glass thickness below 200 μm (‘coverslip’). As a cost-saving and flexible alternative, we recently described COP foils for the fabrication of micropatterned substrates based on a photolithographic approach (Hager et al., 2019).

Here, we present a technological extension of this method for functionalization of COP substrates using large-area polydimethylsiloxane (PDMS)-based elastomeric stamps. The fabrication process of the micropatterned COP foils using large-area μCP is depicted in **Figure 1**. To generate a substrate surface with a high density of oxygen-containing functional groups, such as carboxyl and hydroxyl groups, the COP foil was air-plasma activated in an initial step. Next, a multi-purpose layer of epoxide functional groups was created for subsequent covalent biomolecule binding. The protein patterned cell substrate was finally produced by printing a micrometer-sized BSA grid (for surface passivation) on the epoxysilane-coated COP surface. In order to compensate for the low rigidity of the COP foil, the patterned substrate was bonded with a 384-well plastic casting (**Figure 1A**), resulting in a ready-to-use multi-well plate that can be further functionalized in a modular manner for subsequent analysis of PPIs in live cells with high experimental throughput. **Figure 1B** shows TIRF microscopy images as well as a scanning electron microscope (SEM) recording of a representative 3 μm BSA-Cy5 micropatterned COP surface. The fill-up with Alexa Fluor 488 conjugated streptavidin demonstrates the highly specific binding of proteins in the non-passivated regions. A schematic illustration of the surface chemistry and functionalization with streptavidin and biotinylated antibodies as a basis for subsequent cell experiments is depicted in **Figure 1C**.

**Figure 1.**
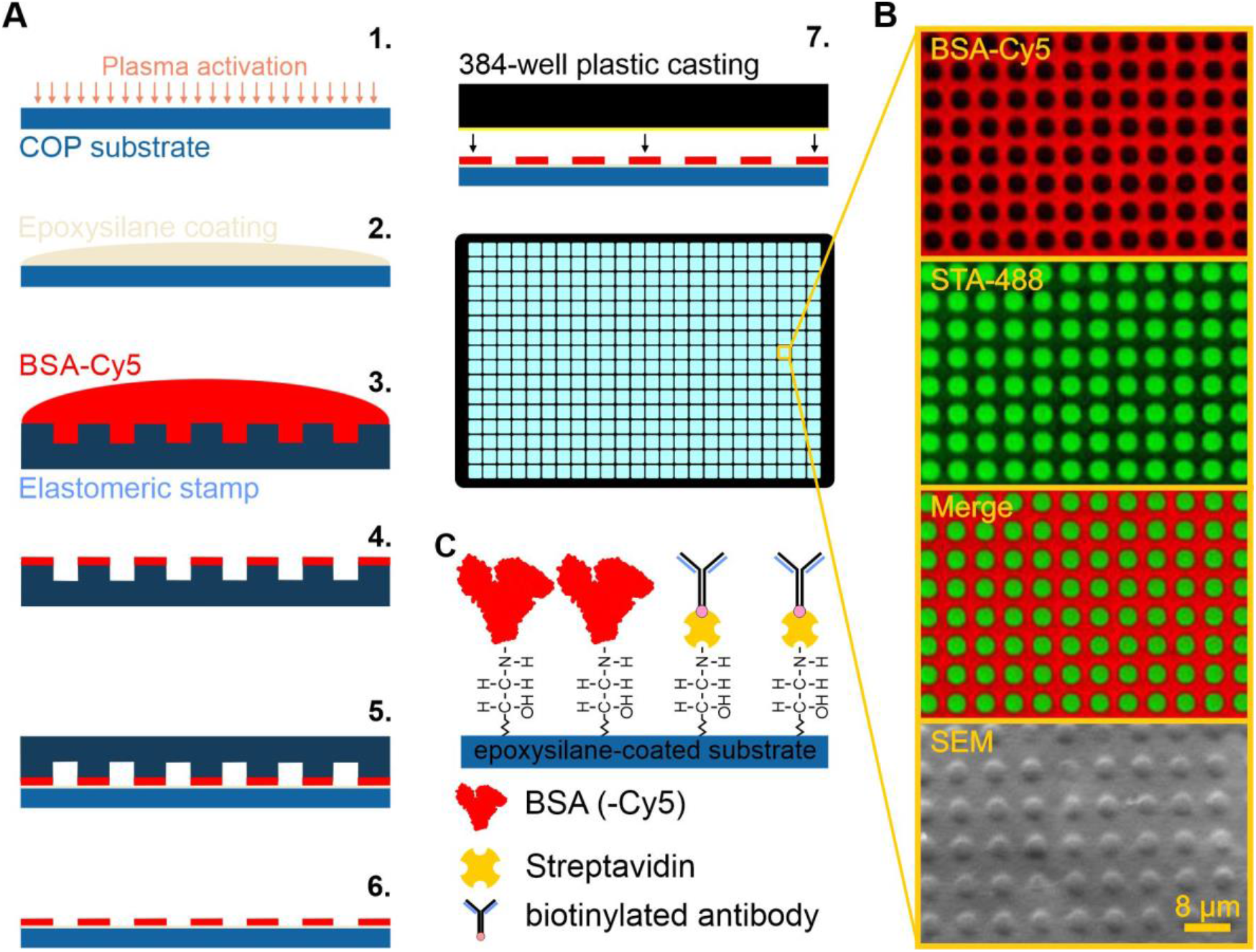
Overview of microcontact printing (μCP) on COP substrates. (A) Schematic workflow of the μCP procedure. In short, COP foils are activated by air-plasma oxidation followed by the introduction of epoxide functional groups. Next, a large-area PDMS stamp containing a continuous grid pattern with a feature size and depth of 3 μm is incubated with a BSA-Cy5 (or BSA) solution for surface passivation. After a washing step, the stamp is placed onto the substrate by its own weight. After stripping-off the stamp, the functionalized COP substrate is bonded with a 384-well plastic casting. (B) Detailed section of the ready-to-use 384-well plate surface for live-cell experiments. Representative TIRF microscopy images of the BSA-Cy5 grid (red, top), Alexa Fluor 488 conjugated streptavidin (green, middle), and merged images (bottom) are shown. (C) Schematic drawing of surface functionalization for live cell experiments consisting of passivated BSA grid, covalent streptavidin-binding in between and addition of biotinylated anti-bait antibodies for specific bait-capturing.

To the best of our knowledge, we demonstrate the first approach for PDMS-based large-area μCP of biomolecules on a COP substrate with micrometer resolution. We are certainly aware that μCP has several methodological limitations, mainly with respect to selectivity (how much protein is adsorbed by the substrate), homogeneity (how much the protein density within the patterned regions varies), and flexibility (each PDMS layout stamp needs a separate mask; and μCP of multiple proteins is difficult due to alignment problems). However, μCP also provides some unexcelled properties compared to other, more sophisticated protein patterning technologies, especially in combination with our high-content micropatterning platform. By use of a customized silicon master (100 mm in diameter) containing a full array of round shaped pillars with a feature size and a depth of 3 μm, we were able to create a large microstructured PDMS stamp for subsequent straightforward functionalization of COP foils. PDMS itself has various advantageous properties for a stamp material: it possesses a hydrophobic surface with low surface energy (favorable for protein transfer onto the target surface), it is chemically inert and elastomeric (molds with high fidelity and can be easily removed from the mask as well as from the substrate), it can be reused, it is cheap and stamping with PDMS is comparatively easy to perform (Alom Ruiz and Chen, 2007; Ricoult et al., 2014; Lindner et al., 2018). Most importantly, our protein patterning approach is robust, highly reproducible, easy to implement (no special and expensive lab equipment is necessary) and tremendously fast (30 min PDMS inking with protein solution and stamping over night followed by bonding step). Additionally, functionalized substrates can be further modified in a modular manner (e.g. with DNA-based systems) (Lanzerstorfer et al., 2020).

### Experimental strategy and optimization for profiling cytosolic protein complexes in the live cell

Based on this antibody patterning approach, we have recently investigated PPIs between various membrane-anchored bait and intracellular prey molecules (Lanzerstorfer et al., 2014; Zindel et al., 2015; Lanzerstorfer et al., 2015; Wagner et al., 2017; Lanzerstorfer et al., 2020). To expand this method for the analysis of exclusively cytosolic PPIs, we adopted the approach of Gandor et al. (2013) and further developed it for investigation of a broadened spectrum of intracellular PPIs with enhanced experimental throughput (**Figure 2**). We therefore combined the use of bait-presenting artificial receptors with our modular and robust large-scale protein patterning platform (**Figure 2A**). The bait-PAR consists of i) a selected intracellular bait protein, ii) an inert transmembrane domain (PDGF receptor transmembrane domain), and iii) a flexible extracellular domain (four repeats of Titin Ig I27 domain) that contains a human influenza hemagglutinin (HA) epitope tag, which directs the artificial receptor towards the patterned anti-HA antibodies (**Figure 2B**). The bait-PAR as well as the cytosolic prey is expressed as a fluorescent fusion protein and PPIs are monitored by the degree of bait-prey copatterning using TIRF microscopy. In order to exemplify the validation and broad applicability of this assay, we constructed a bait-PAR consisting of the growth factor receptor-bound protein 2 (Grb2), herein referred to as bait-PAR-Grb2. Grb2 is a widely expressed cytosolic adapter protein and acts as an intermediate between cell-surface activated receptors and downstream targets through SH2 and SH3 domains (Pawson, 2007). Furthermore, Grb2 is reported to mediate intracellular signalling dynamics by the interaction with a variety of downstream molecules (Bisson et al., 2011), and therefore represents a perfect intracellular proof of concept target. In a first attempt, the correct bait-PAR-Grb2 orientation across the cell membrane as well as the cell membrane-substrate interface was investigated as prerequisites for further analysis using TIRF microscopy. For this purpose, GFP-fused bait-PAR-Grb2 was transiently expressed in Hela cells, which were subsequently incubated on an anti-HA antibody patterned COP substrate (**Figure 2C**). For quantitation of the lateral bait and prey distribution, the respective fluorescence signal intensities within and outside the antibody-patterned areas were compared. The fluorescence contrast (signal ratio) provides a measure for the specificity of bait enrichment as well as for the bait-prey interaction strength (Schütz et al., 2017). Bait-PAR-Grb2 was found to be significantly enriched in the cognate antibody-functionalized micropatterns (mean fluorescence contrast <c> 0.38 ± 0.02), indicating a correct integration of the artificial construct into the plasma membrane as well as a high specificity of antigen-antibody binding. On the contrary, we observed a homogenous staining with the lipophilic dye DiD (mean fluorescence contrast <c> 0.09 ± 0.01), proofing a flat interface of the cell membrane with the patterned COP substrate to avoid false positive results. As Hela cells were shown to fulfil those requirements, they were used throughout the study. Compared to the control conditions, we detected a ~4-fold increase in bait protein enrichment in antibody patterned areas. This value is comparable to other studies using different protein patterning approaches (Singhai et al., 2014; Dirscherl et al., 2018; Watson et al., 2021), again proofing the effective surface functionalization of our platform.

**Figure 2.**
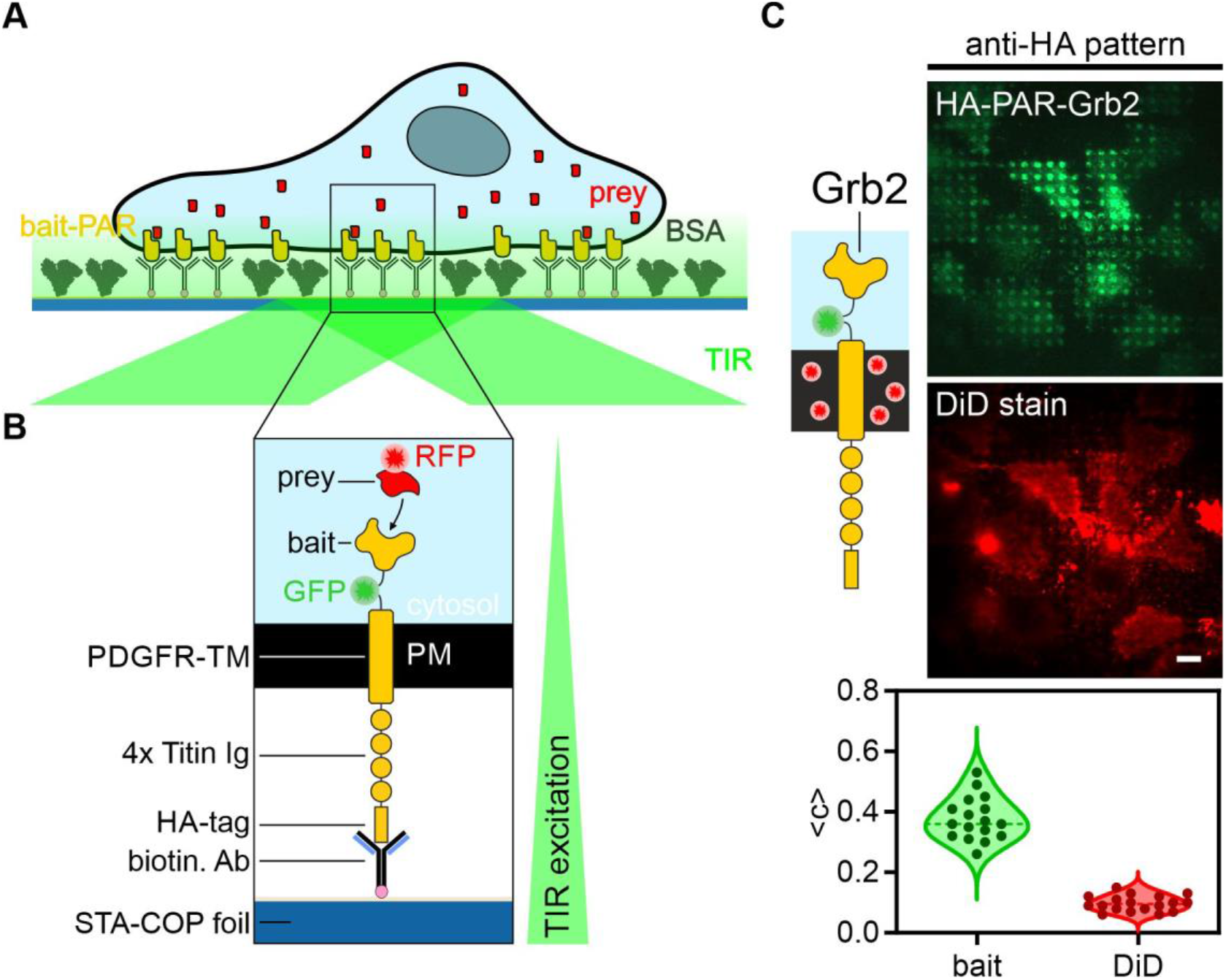
Patterning of bait-presenting artificial receptors (bait-PARs) for visual coimmunoprecipitation of cytosolic protein complexes. (A) Schematic presentation of the micropatterning assay. Cells are transiently co-transfected with bait-PARs fused to GFP (or RFP) and RFP-labeled (or GFP) prey molecules. Upon specific antibody-antigen interaction, bait-PARs are rearranged in the plasma membrane according to the micrometer-scale antibody pattern on the COP substrate. The interaction between bait-PARs and the prey is monitored by the degree of prey copatterning. (B) Schematic illustration of a single bait-PAR. The bait-PAR is composed of an intracellular bait protein, a conjugated fluorophore, a single transmembrane domain, an extracellular spacer domain (four repeats of the Titin Ig domain I27) and a HA epitope tag, which directs the bait-PAR towards the pattern of the cognate immobilized anti-HA antibody. (C) Adaption of the bait-PAR assay for analysis of cytosolic protein complexes downstream the EGFR. The bait protein (regulatory subunit of protein kinase A) of the previously published bait-PAR (Gandor et al., 2013) was exchanged with the growth-factor receptor binding protein 2 (Grb2). In order to proof sufficient cell attachment to micropatterned COP substrates as a prerequisite for TIRF microscopy, Hela cells were transiently transfected with HA-PAR-Grb2 and cell membrane was stained with the lipophilic tracer DiD. Scale bar: 9 μm. Abbreviations: biotin. Ab, biotinylated antibody; HA-tag, human influenza hemagglutinin epitope tag; PDGFR-TM, transmembrane domain of PDGF receptor; PM, plasma membrane; STA-COP foil, streptavidin-coated COP foil.

In a next step, the functionality of Grb2 fused to the bait-PAR was tested (**Figure 3**). Therefore, we used the well-described interaction between the EGFR and Grb2. As Grb2 has been reported to directly bind phosphotyrosine (pTyr)-containing sequences on the EGFR via its SH2 domain (Yamazaki et al., 2002; Jiang et al., 2003), we used the EGFR as the bait and Grb2 coupled to the artificial receptor construct as the prey (**Figure 3A**). Cells expressing GFP-labeled bait-PAR-Grb2 were grown on an anti-EGFR patterned surface and were imaged using TIRF microscopy before and after EGF stimulation. The degree of bait-PAR-Grb2 copatterning to EGFR-enriched areas served as a parameter of EGFR downstream signalling activation. Under basal conditions, bait-PAR-Grb2 showed minor colocalization with EGFR enriched areas, whereas a significant copatterning was detected upon EGF stimulation within minutes, indicating that Grb2, despite coupled to the artificial transmembrane domain, can still translocate and bind to the ligand-activated EGFR.

**Figure 3.**
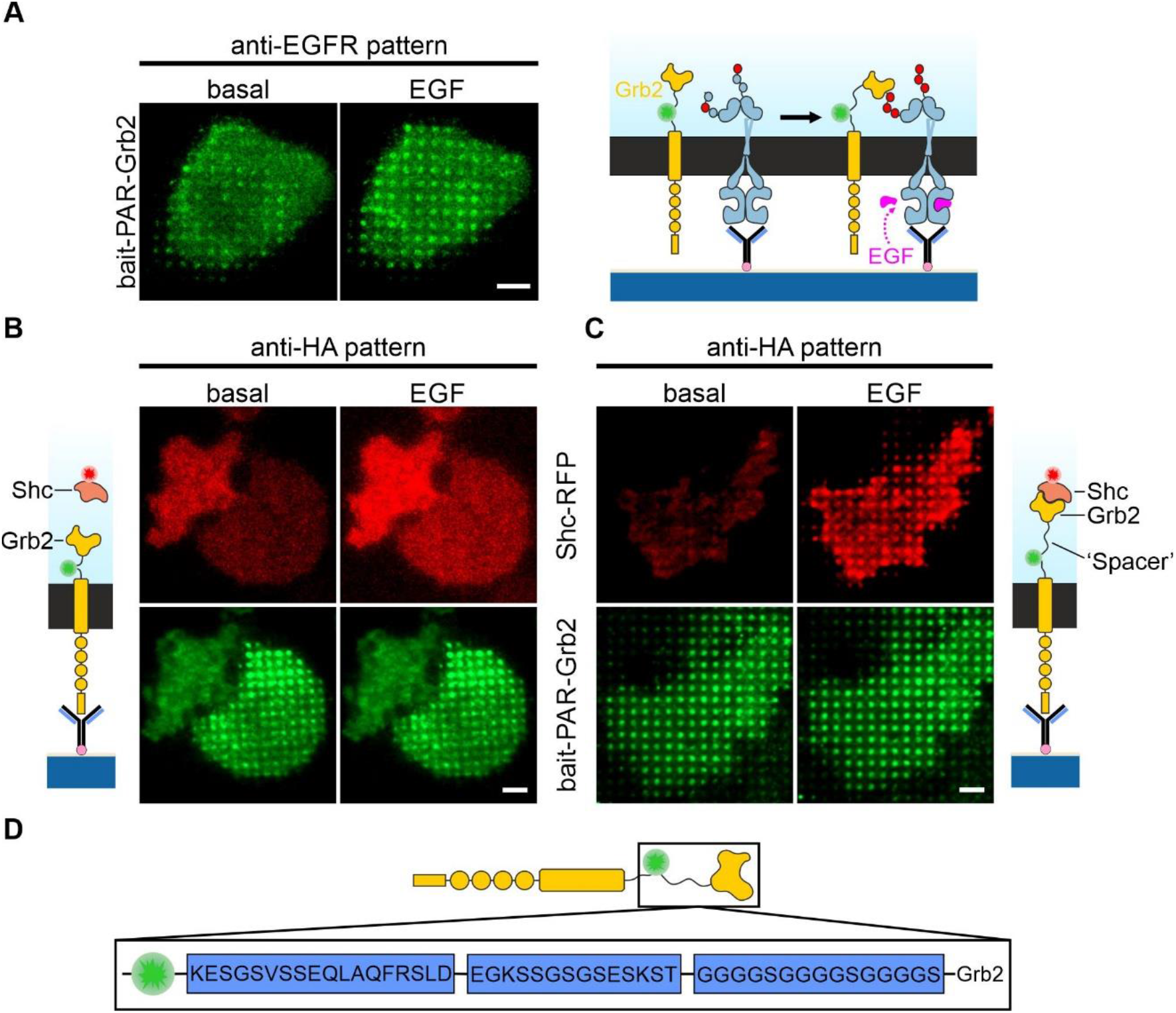
Test of bait functionality coupled to artificial receptor construct and optimization for protein complex formation. (A) Cells expressing modified bait-PAR consisting of wildtype Grb2 were grown on anti-EGFR patterned surfaces. Copatterning of bait-PAR-Grb2 to EGFR-enriched micropatterns was analyzed before and after EGF stimulation (170 nM, 10 min). Scale bar: 15 μm. (B) Cells co-expressing bait-PAR-Grb2 (GFP-fused, green) and Shc-RFP (RFP-fused, red) were grown on anti-HA patterns and copatterning of Shc to bait-PARs upon EGF stimulation (170 nM, 10 min) was assessed by TIRF microscopy. Representative images of cells expressing adapted bait-PAR-Grb2 (B) and optimized bait-PAR-Grb2 with additional amino acid sequences to enhance flexibility of the bait protein (C). Scale bar: 12 μm. (D) Schematic drawing of optimized bait-PAR-Grb2 with inserted linker sequences between fluorophore and bait protein.

To proof Grb2 activation and mediation of downstream signalling, we next investigated the ability of bait-PAR-Grb2 to corecruit further adapter proteins such as the SHC-transforming protein 1 (Shc1), which has been reported to be recognized by the Grb2 SH2 domain (Salcini et al., 1994), similar to the EGFR. Therefore, cells coexpressing bait-PAR-Grb2 (GFP-fused) and Shc1-RFP were grown on anti-HA patterned surfaces and Shc1 copatterning was monitored upon EGF stimulation (**Figure 3B**). Surprisingly, we could not detect any significant Shc1 colocalization to the bait-PAR-Grb2 patterned areas, neither in unstimulated nor in EGF stimulated cells. However, we found a substantial RFP fluorescence intensity increase under TIR illumination conditions upon EGF addition, indicating an agonist dependent Shc1 translocation to the cell membrane. To further elaborate on this issue, we intended to enhance the flexibility of the intracellular portion of the bait-PAR by inserting flexible fusion protein linkers between the fluorophore and Grb2 (**Figure 3D**) (Chen et al., 2013). Indeed, in cells coexpressing the optimized bait-PAR-Grb2 and Shc1-RFP, a prominent Shc1 copatterning was detected upon EGFR activation, indicating Grb2-Shc1 protein complex formation (**Figure 3C**). We therefore used the optimized and more flexible bait-PAR-Grb2 for subsequent experiments. A similar linker system was recently reported by Incaviglia et al. (2020), investigating the Grb2:SOS1 complex by focal molography.

Protein micropatterning can lead to the formation of protein clusters within or at the cell membrane with subsequent recruitment of relevant proteins (Watson et al., 2021), including bait and prey molecules of interest (Lanzerstorfer et al., 2020). To investigate unspecific bait-prey copatterning in the presented approach, bait-PAR and prey distribution was checked on microstructured surfaces but without antibody incubation (**Figure S1**). Neither under basal conditions, nor after EGF stimulation, an unspecific copatterning was detected for all bait and prey proteins under study. We therefore conclude that bait-prey copatterning occurs due to interactions and active corecruitment.

### Visual immunoprecipitation reveals differences in Grb2-mediated protein assemblies downstream the EGFR

The EGFR is a tyrosine kinase and is found to be upregulated in different types of cancers, mainly caused by mutations and truncations of its extracellular as well as its intracellular kinase domain. Consequently, the two main pro-oncogenic downstream signalling pathways, the Ras-Raf-MEK and PI3K-Akt pathway, are frequently over-activated (Wee and Wang, 2017). Hence, it is of critical importance to understand the molecular mechanisms that regulate EGFR signal transduction. Within this regard, cytosolic proteins downstream the EGFR are attracting notice as key regulatory targets, particularly Grb2, as it is one of the most important proteins participating in EGFR signalling. Grb2 serves as an universal adapter protein once the EGFR is activated, subsequently leading to the activation of the aforementioned pro-oncogenic signalling pathways (Yin et al., 2013). To analyze Grb2-mediated protein complexes with high fidelity in a live cell context, we aimed in the visual immunoprecipitation of protein assemblies within the Ras-Raf-MEK and PI3K-Akt pathway by use of the bait-PAR system (**Figure 4 and 5**).

**Figure 4.**
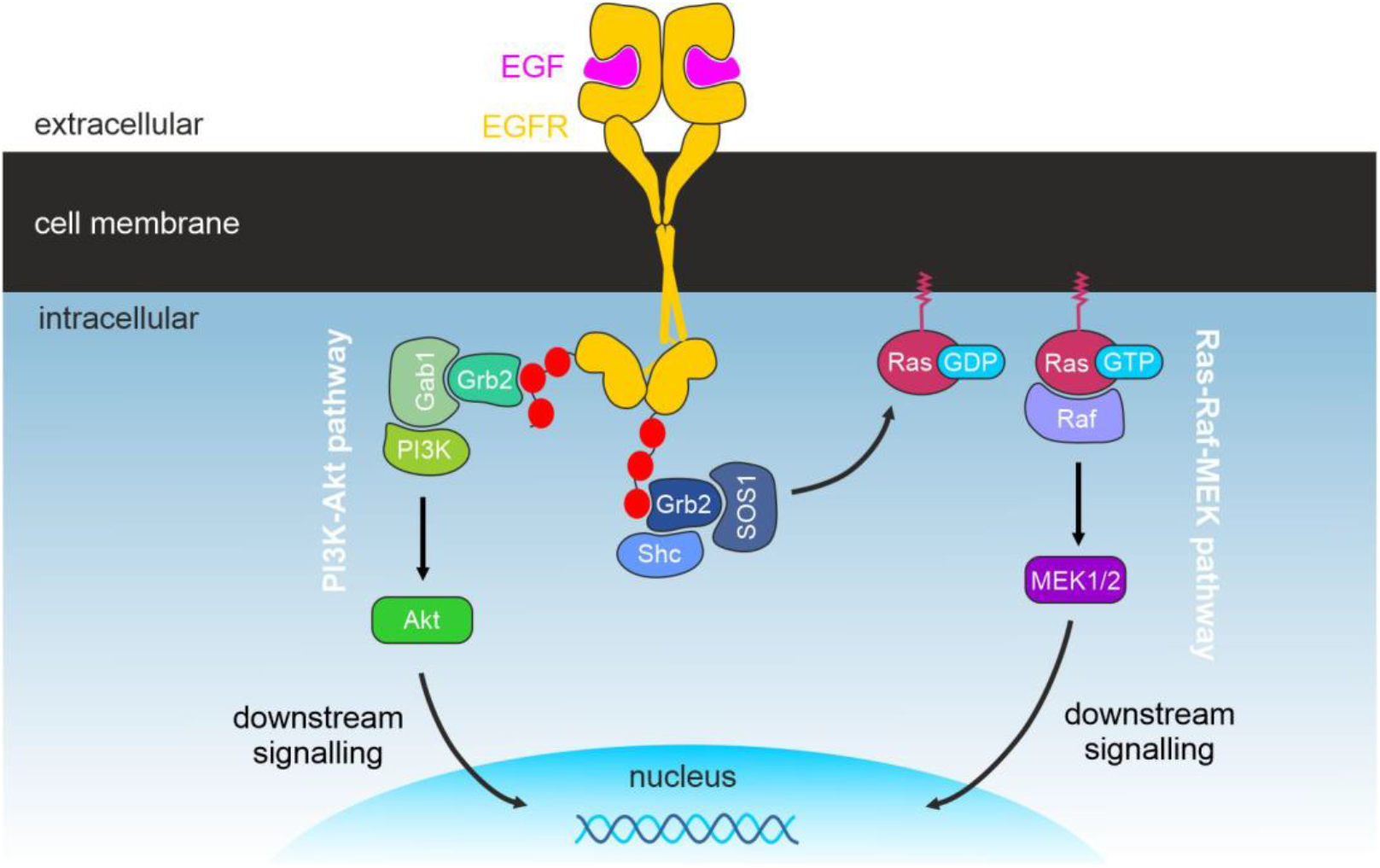
Illustration of the two main pro-oncogenic EGFR downstream signalling cascades and ligand-induced assembly of protein complexes. Depicted are the Ras-Raf-MEK and the PI3K-Akt pathways.

The Ras-Raf-MEK signal transduction pathway is initiated by EGFR activation through binding of its cognate ligands (epidermal growth factor (EGF) and transforming growth factor α (TGF α)), leading to EGFR dimerization and activation of its cytoplasmic tyrosine kinase domain (Tomas et al., 2014). Subsequently, a ternary complex consisting of Shc1:Grb2:SOS1 (son of sevenless protein 1) is recruited to the phosphorylated RTK (Arkun and Yasemi, 2018), which further leads to the activation of the membrane-bound small GTPase protein Ras (rat sarcoma protein). Upon exchanging GDP for GTP, Ras in turn activates the serine/threonine-specific protein kinase Raf (rapidly accelerated fibrosarcoma protein), leading to sequential phosphorylation and activation of the respective downstream signalling cascade (Wennerberg et al., 2005).

We first investigated the initial protein complex formation within the Ras-Raf-MEK pathway between Grb2, SOS1 and Shc1. Grb2 is known to be constitutively bound to SOS1, predominantly via its N-terminal SH3 domain (Yu et al., 2017), whereas Shc1 associates with Grb2 upon EGFR stimulation via the SH2 domain (Ravichandran et al., 1995) (**Figure 5A-D**). In cells coexpressing RFP-fused bait-PAR-Grb2 and SOS1-GFP, we indeed found a prominent SOS1 copatterning to bait-enriched micropatterns under basal conditions (<c_prey/bait_> 0.55 ± 0.03), which did not change upon EGF stimulation (<c_prey/bait_> 0.56 ± 0.02), again indicating an agonist-independent stable association between Grb2 and SOS1 (**Figure 5A and C**). On the contrary, in cells coexpressing GFP-fused bait-PAR-Grb2 and Shc1-RFP, we could confirm the agonist-dependent Grb2:Shc1 complex formation as indicated by a low degree of copatterning in unstimulated cells (<c_prey/bait_> 0.14 ± 0.02), and a significant increase in Shc1 corecruitment (*p < 0.0001*) upon EGF stimulation (<c_prey/bait_> 0.45 ± 0.03) (**Figure 5B and C**).

**Figure 5.**
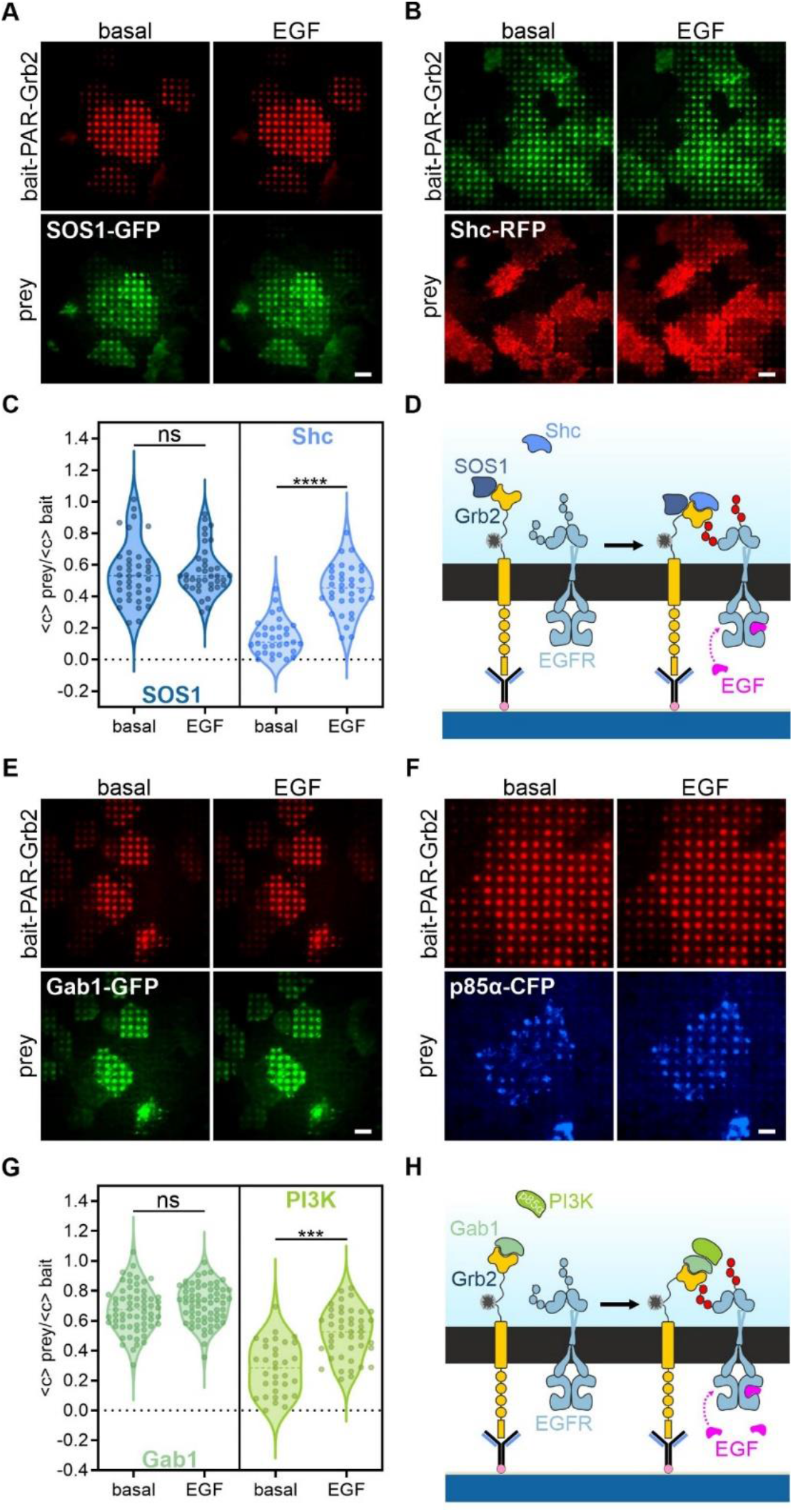
Visual immunoprecipitation reveals differences in EGFR-mediated cytosolic protein complexes. Initial signalling complexes of Ras-Raf-MEK (A-D) and PI3K-Akt pathway (E-H). Hela cells were transiently co-transfected as the following: (A) bait-PAR-Grb2-RFP + SOS1-GFP, (B) bait-PAR-Grb2-GFP + Shc-RFP, (E) bait-PAR-Grb2-RFP + Gab1-GFP, and (F) bait-PAR-Grb2-RFP + p85α-CFP. Transfected cells were grown for at least four hours on anti-HA antibody patterned substrates 24 hours after transfection. Shown are representative TIRF microscopy images of cells expressing fluorescently labelled bait and prey proteins before and after EGF stimulation for 10 min (170 nM) (A, B, E, F). Scale bar: 9 μm. Schematic presentations illustrate indicated protein complex assembly (D, H). Charts show quantitation of bait-normalized fluorescence contrast of respective prey copatterning before and after EGF addition (C, G). Error bars are based on mean ± SE of at least 35 analyzed cells measured on three different days. *** p < 0.001 and **** p < 0.0001 for comparison of bait-normalized prey copatterning before and after EGF stimulation. ns, no significant differences.

Besides the Ras-Raf-MEK signalling, the PI3K-Akt pathway is the second major EGFR-mediated signal transduction pathway, involving binding of Grb2 to the EGFR and subsequent association with Gab1 (Grb2-associated-binding protein 1; predominantly via the C-terminal SH3 domain) and the p85 subunit of PI3K (phosphoinositide 3-kinases; via pTyr residues of Gab1), resulting in the production of phosphatidylinositol (3,4,5)-triphosphate (PIP3) and activation of Akt (Wee and Wang, 2017). Like SOS1, Gab1 forms a constitutive complex with Grb2 (Schaeper et al., 2000), whereas the association between Grb2:Gab1 and PI3K-p85 can be enhanced by EGF addition (Holgado-Madruga et al., 1996) (**Figure 5E-H**). Again, we could detect a constitutive Grb2:Gab1 complex formation in cells coexpressing RFP-fused bait-PAR-Grb2 and Gab1-GFP, as indicated by the prominent Gab1 copatterning under basal conditions (<c_prey/bait_> 0.67 ± 0.02) (**Figure 5E and G**). Similar to SOS1, the Gab1 fluorescence contrast did not change significantly upon EGFR activation (<c_prey/bait_> 0.71 ± 0.02). For the investigation of PI3K association, we coexpressed the CFP-fused p85α regulatory subunit of PI3K and analyzed the copatterning to bait-PAR-Grb2 (**Figure 5F and G**). The Grb2:Gab1 complex readily showed an association with p85α in the absence of growth factor (<c_prey/bait_> 0.28 ± 0.03), but this interaction was significantly enhanced by EGF addition (*p < 0.001,* <c_prey/bait_> 0.51 ± 0.02), suggesting further interaction between pTyr residues of Gab1 and p85α.

We next questioned whether those ternary protein complexes can actively recruit further downstream molecules. Therefore, the subcellular localization of proximate scaffold proteins such as Ras, Raf and MEK1 (mitogen-activated protein kinase kinase 1) was investigated (**Figure S2**). Ras is activated by the guanine nucleotide exchange factor SOS1 by induction of the exchange of GDP to GTP (Wee and Wang, 2017). So far it is not clear whether this occurs through dissociation of the Grb2:SOS1 complex from the receptor and translocation to the membrane-bound Ras, or by active recruitment of Ras to the activated Grb2:SOS1 complex. As shown in **Figure S2A**, Ras can be actively copatterned to bait-PAR-Grb2 enriched areas, however, the majority of analyzed cells showed a homogenous HRas-CFP membrane distribution, indicating that Grb2:SOS1 or SOS1 alone dissociates from the receptor complex to activate Ras at the plasma membrane. Furthermore, a Grb2-independent SOS1 membrane-localization and receptor-triggered Ras activation has been recently reported (Christensen et al., 2016), which could also explain our observation. Additionally, we cannot fully exclude a reduced SOS1:Ras interaction caused by spatial restrictions due to the artificially patterned Grb2:SOS1 complex. However, a similar appearance was also obtained for the downstream effector Raf, which is subsequently corecruited and activated by Ras (Wee and Wang, 2017). Raf1-CFP copatterning to bait-PAR-Grb2 patterns was a rather rare event, as in most of the cells Raf1 showed a homogenous membrane recruitment upon EGF stimulation, independently of bait-PAR-Grb2 micropatterns (**Figure S2B**). No copatterning was detected for MEK1-GFP, which in turn is activated by Raf (**Figure S2C**).

Taken together, we could clearly show, that our visual immunoprecipitation assay is suitable to generally characterize cytosolic protein complex formation. Moreover, we were able to confirm and to discriminate between constitutive protein complexes and agonist induced associations, which were mainly investigated by classical biochemical approaches such as coimmunoprecipitation in the past.

### Modulation of cytosolic PPIs by protein complex disruptors

Recent studies evidenced that Grb2 is involved in the development and progression of multiple tumor malignancies such as breast, lung and bladder cancer, chronic myelogenous leukemia, hepatocellular carcinoma, etc. (Ijaz et al., 2018). Therefore, Grb2 has become an attractive therapeutic target, mainly by modulating its downstream signalling activity by peptidomimetics via blocking its SH2 (connection to cell surface receptors via Shc1 interaction) and SH3 (interlink to downstream pathways) domains (Gao et al., 2000; Burke, 2006; Gril et al., 2007; Morlacchi et al., 2014). To demonstrate the applicability of our assay to study PPI inhibitors in living cells, we monitored the dissociation behaviour of the constitutively bound SOS1: Grb2:Gab1 ternary signalling complex (**Figure 6**).

**Figure 6.**
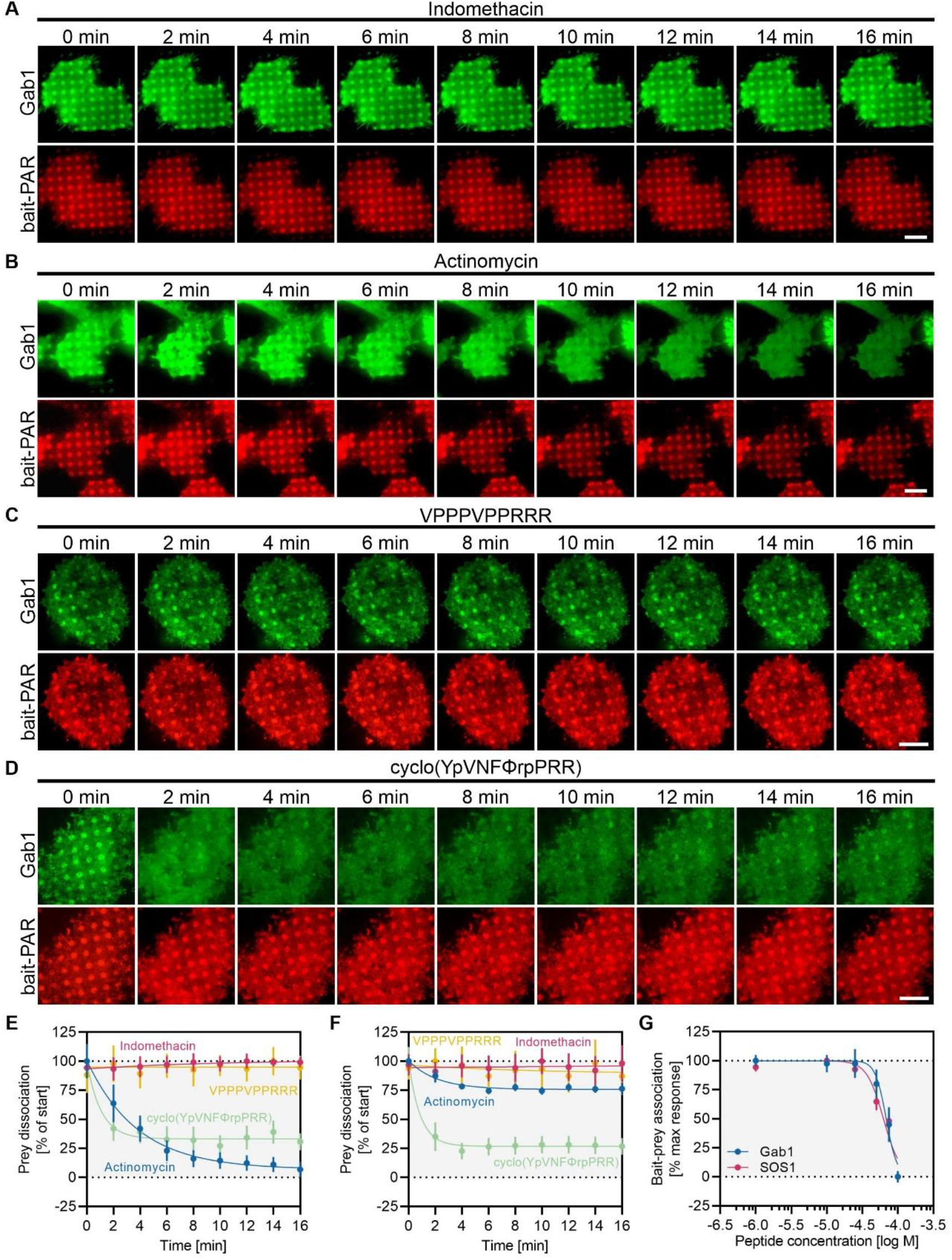
Disruption of protein complexes by protein domain inhibitors. Cells co-expressing bait-PAR-Grb2-RFP and Gab1 – GFP were used to showcase the different effects of indicated inhibitory substances (A) – (D). Representative TIRF microscopy images show co-recruitment of the prey to bait micropatterns before and after pharmacological treatment. Scale bars: 15 μm. Quantitation of bait-normalized fluorescence contrast of Gab1 (E) and SOS1 (F) dissociation kinetics upon substance treatment. (G) Dose-response relationship of cell-permeable disruptive peptide. Data represent mean ± SE of > 40 analyzed cells measured on at least two different days.

To this end, Hela cells expressing bait-PAR-Grb2-RFP and GFP-fused SOS1 or Gab1 were stimulated with various reported disruptive substances and bait-prey copatterning was monitored over time. To showcase the different effects of agents under study (10 μM indomethacin, 10 μM actinomycin D, 100 μM peptide VPPPVPPRRR and 100 μM peptide cyclo(YpVNFΦrpPRR)), representative TIRF microscopy images of cells co-expressing bait-PAR-Grb2-RFP and Gab1-GFP are depicted in Figure 6A-D. Indomethacin, a known nonsteroidal anti-inflammatory drug (NSAID) and recently identified inhibitor of the Shc1:EGFR interaction (Lin et al., 2019), was used as a negative control for the N- and C-terminal SH3 domain mediated SOS1:Grb2:Gab1 interaction. No effect on Gab1 (Figure 6A and E) and SOS1 (Figure 6F) copatterning was detected over a time period of 16 min. On the contrary, actinomycin D, a reported anti-cancer drug and Grb2 SH2 domain inhibitor (Kim et al., 2000), was identified also as a potent SH3 domain inhibitor, resulting in a gradual dissociation of Gab1 (Figure 6B and E) and SOS1 (Figure 6F) from Grb2. For Gab1, copatterning was remarkably reduced by ~95% with a dissociation half-life of 2.6 min, whereas SOS1 copatterning was reduced by ~25% (dissociation half-life of 1.4 min), indicating a more pronounced affinity of actinomycin D for the C-terminal SH3 domain of Grb2, which mediates binding of Gab1. In recent years, high affinity Grb2-binding peptides have been developed to block Grb2 association to cell surface receptors (Chen et al., 2010; Noguchi et al., 2017) or binding to downstream molecules (Kardinal et al., 2001; Gril et al., 2007). We therefore further tested two known Grb2 inhibitors, the SH3 domain blocking peptide VPPPVPPRRR (Vidal et al., 1999) and the most recently described Grb2 SH2 domain inhibitor cyclo(YpVNFΦrpPRR) (Wen et al., 2020). Upon stimulation with 100 μM VPPPVPPRRR we could not detect any effect on Gab1 and SOS1 copatterning (Figure 6C, E and F). This observation might be readily explained by a general low lipid membrane permeability of peptides (Yang and Hinner, 2015). Thus, extensive effort has been made to develop novel cyclic cell-penetrating peptides (CPPs), that are in addition capable of binding to target proteins with antibody-like affinity and specificity, such as the CPP cyclo(YpVNFΦrpPRR) (Wen et al., 2020). Indeed, when cells were treated with 100 μM cyclo(YpVNFΦrpPRR), we observed a rapid dissociation of Gab1 (Figure 6D and E) and SOS1 (Figure 6F) from Grb2, reaching a maximum prey dissociation of 70-75% already after 4-6 minutes of peptide treatment. The comparable dissociation half-lifes of 0.6 min (SOS1) and 0.7 min (Gab1) indicate a similar affinity of cyclo(YpVNFΦrpPRR) for the N- and C-terminal SH3 domains of Grb2. Our results indicate that the peptide cyclo(YpVNFΦrpPRR) does not only block the Grb2 SH2 domain as previously reported (Wen et al., 2020), but also the SH3 domains, which indicates that the assay has the ability of identifying novel inhibitory targets. Peptide cyclo(YpVNFΦrpPRR) was reported to dose-dependently reduce the level of phosphorylated MEK (p-MEK) with an IC50 value of ~ 15 μM. Therefore, we further elaborated on the half-maximal effective peptide concentration (EC50), which is necessary to dissolve the bait-prey interaction in our system (Figure 6G). In line with the comparable dissociation properties, we observed similar EC50 values for both prey proteins, with 68 μM for Gab1 and 63 μM for SOS1. The ~4-fold increase in peptide concentration compared to the reported value of 15 μM might be presumably caused by a lower affinity and blocking efficacy of SH3 domains in comparison to the SH2 domain.

Altogether, we could demonstrate that the assay is capable of determining putative differences in the specificity, efficacy and affinity of known as well as unknown protein domain inhibitors.

### Monitoring protein complex formation dynamics in individual cells

It is now obvious that distinct PPI dynamics such as interaction lifetime, binding affinity, and protein complex stability are important regulators of fundamental processes in living cells. Therefore, the spatiotemporal manipulation and monitoring of signalling events is key to interlink the nature of dynamic signalling and its importance for information transfer and cell response (Doupé and Perrimon, 2014). In order to learn how the cytosolic environment in a cell impacts protein complex formation and signalling rates, it is of particular importance to perform measurements in living cells rather than doing biochemical analysis in dilute solutions. Moreover, it is also appreciated to perform measurements on a single cell level to unravel cell-to-cell heterogeneities, as even genetically identical cells can behave differently (Ryu et al., 2019).

As shown in previous studies, the micropatterning approach is a superior tool to study protein interaction kinetics in a live cell context (Schwarzenbacher et al., 2008; Weghuber et al., 2010; Löchte et al., 2014; Lanzerstorfer et al., 2015; Zindel et al., 2015; Motsch et al., 2019). To monitor protein complex formation dynamics in individual cells, we carried out TIR-based fluorescence recovery after photobleaching (TIR-FRAP) experiments (**Figure 7**). We therefore used this approach to further elucidate on the different observed interaction regimes. For this purpose, cells cotransfected with bait-PAR-Grb2 and different prey molecules were grown on anti-HA antibody patterned surfaces and single patterns were bleached using a high-intensity laser pulse for the determination of the temporal prey fluorescence recovery dynamics (**Figure 7A**). **Figure 7B** shows the respective fluorescence recovery curves for the indicated prey proteins. Depending on their lifetime, PPIs can be discriminated into permanent or transient interactions, whereas the latter ones are crucial for short-lived biological processes such as signal transduction (Wang et al., 2014). In general, the recovery process of the three investigated prey molecules (Shc1, SOS1, Gab1) proved to be fast, indicating transient PPIs. From the FRAP curves, the exchange rate of the freely diffusing pool of prey molecules into and out of the bleached ROIs was obtained through a bi-exponential fit as the slow recovery rate (k_slow_) (**Figure 7E**), whereas the fast recovery rate (k_fast_) represents free diffusion (Sprague and McNally, 2005). A biphasic binding behaviour of Grb2 to adapter proteins was previously reported (Mol et al., 2004). In living cells, such a two-step model could be described with an initial diffusion step of the adapter protein (here the prey protein) from cytosolic compartments to the membrane interface, followed by a second step including specific bait-prey binding/rebinding events. Interestingly, Gab1 and SOS1, which were found to be constitutively bound to Grb2, exhibited a significantly lower exchange rate (k_slow_) than Shc1, which was shown to interact with Grb2 in an agonist-dependent manner (Gab1: 0.030 ± 0.002 s^-1^, SOS1: 0.074 ± 0.004 s^-1^, and Shc1: 0.103 ± 0.003 s^-1^). Those results suggest halftimes of dissociation from the pattern-bound immobile bait-prey associations of about 30 sec for Gab1, 13 sec for SOS1 and 10 sec for Shc1. In the FRAP experiments, a portion of the prey molecules appeared immobile on a timescale of seconds as evidenced by the incomplete fluorescence recovery (**Figure 7D**). In line with the observation for different exchange rates, Gab1 and SOS1 showed significantly decreased mobile fractions (37.8 ± 2.1% and 46.7 ± 1.1%) when compared to Shc1 (55.8 ± 1.5%). The decreased exchange and mobile fraction of Gab1 and SOS1 molecules associated to the patterned Grb2 suggests either multiple association and dissociation evens due to densely immobilized binding partners or indicates a more stable bait-prey association and protein complex stability. As the prey expression level might influence the recovery rates of the bleached molecules, cells with comparable prey expression were used for FRAP experiments (**Figure 7C**).

**Figure 7.**
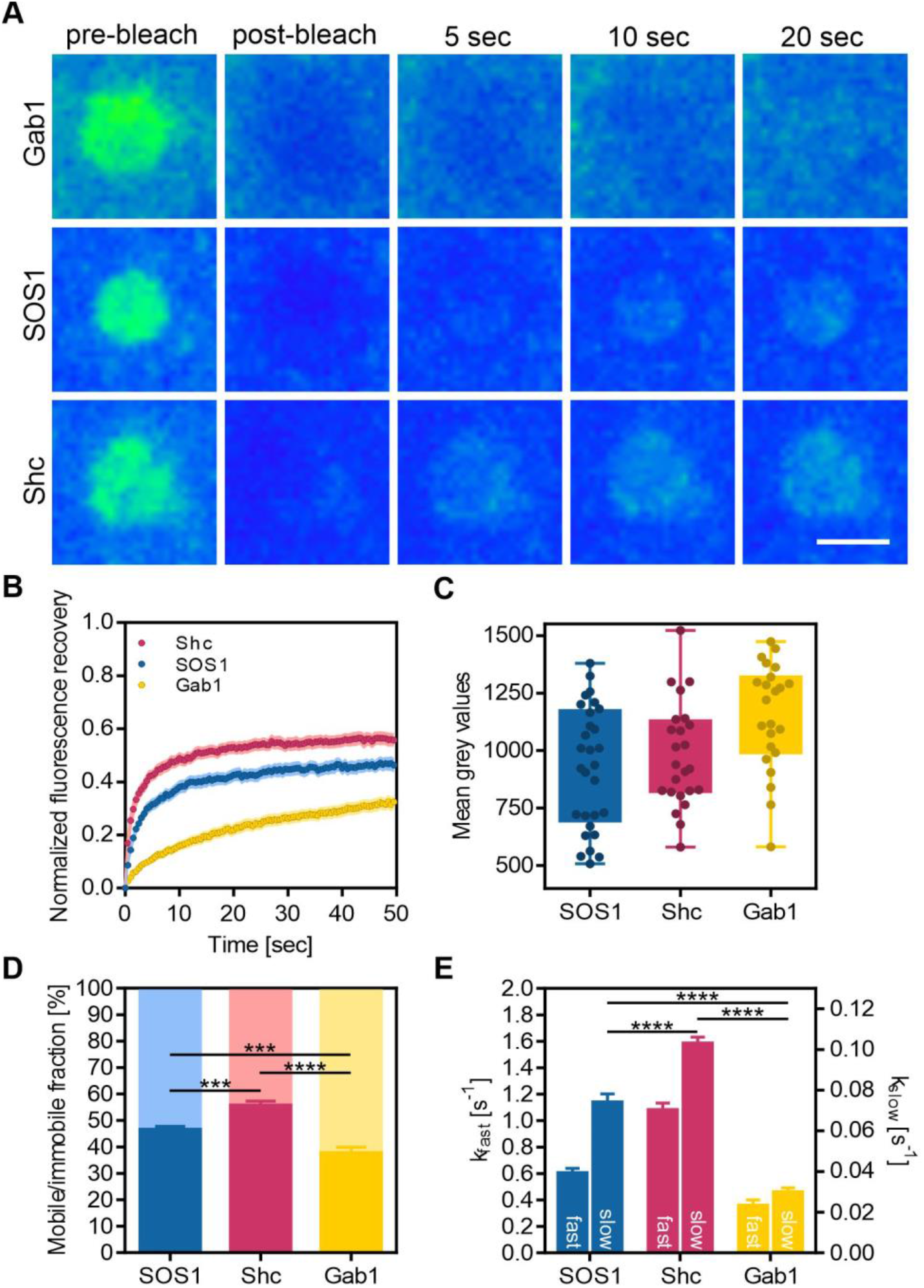
Monitoring protein complex formation dynamics in individual cells. (A) Representative TIR-FRAP images of single bleached prey patterns at indicated time points are shown. For FRAP experiments, cells were co-transfected with bait-PAR-Grb2 and indicated prey proteins and were grown on anti-HA patterned substrates. Prior to FRAP, cells were stimulated with EGF (170 nM) for at least 5 min. Individual patterns were selected for the FRAP experiment. Scale bar: 3 μm. Images shown were intensity adjusted and false coloured for better visualization of differences in prey recovery dynamics. (B) Normalized mean fluorescence recovery curves of analyzed prey molecules. (C) Mean fluorescence intensities of cells used for photobleaching experiments. (D) Calculation of mobile/immobile fraction. *** p < 0.001 and **** p < 0.0001 for comparison of mobile fractions. (E) Calculation of exchange rates of prey molecules from fluorescence recovery curves. Diffusion of prey molecules is represented by k_fast_ (left y-axis). Prey binding to and dissociation from Grb2 is depicted as k_slow_ (right y-axis). **** p < 0.0001 for comparison of kslow. Error bars are based on mean ± SE of at least 25 analyzed cells measured on three different days.

Whereas spatiotemporal modelling and characterization of binding events of cytosolic proteins to membrane receptors, and more precisely to the EGFR, are well described (Hsieh et al., 2010), respective kinetic information on cytosolic PPIs in living cells is missing. We here provide evidence that the temporal regulation of EGFR signalling networks is not solely regulated by unique recruitment and binding signatures of different scaffold proteins to the receptor. Much more it appears that these signalling processes are also defined through the interaction properties of the downstream molecules itself.

## Conclusion

The approach presented in this paper will allow for visual immunoprecipitation of many cytosolic proteins of interest which are at least able to be located to or near the cell membrane interface by use of the bait-PAR construct. Analysis of proteins from intracellular locations other than the cytosol might require further adaption of the bait-PAR with respect to flexibility and range of the cytosolic domain. The subcellular relocalization of signalling molecules in spatially defined micropatterns within single cells enables for in-depth investigation of intracellular protein complexes in its native environment with high specificity using TIRF or confocal microscopy. Furthermore, a more structural characterization might be achieved when combining our subcellular micropatterning assay with high-resolution microscopy techniques such as single molecule microscopy (Löchte et al., 2014; Sevcsik et al., 2015) or CryoEM (Engel et al., 2019; Toro-Nahuelpan et al., 2020). In addition, upon optimization of bait and prey fluorophore positions, resonance energy transfer-based experiments such as FRET, BRET, FLIM and simultaneous FRAP and FRET are possible, which would enable a direct proof of PPIs.

In conclusion, we envision that the method described in this paper will become a valuable alternative or add-on to standard wet lab-based technologies such as biochemical immunoprecipitation.

## Materials and Methods

### Reagents, materials and DNA constructs

Bovine serum albumin (BSA), Tyrphostin AG 1478, actinomycin D, indomethacin, streptavidin, EGF, (3-Glycidyloxypropyl)trimethoxysilane (GPS) (98%), Polydimethylsiloxane (PDMS, SYLGARD^®^ 184) and DMSO were purchased from Sigma-Aldrich (Schnellendorf, Germany). BSA-Cy5 was obtained from Protein Mods (Madison, Wisconsin, USA). Biotinylated anti-EGFR, anti-HA, mouse-IgG and anti-mouse-IgG (FITC) were purchased from Antibodies Online (Herford, Germany). The cell-permeable cyclic peptide cyclo(YpVNFΦrpPRR) was custom synthesized by BioCat GmbH (Heidelberg, Germany) and the VPPPVPPRRR peptide was purchased from Santa Cruz Biotechnology (Dallas, Texas, USA). Cyclic olefin polymer (Zeonor-COP) foils with a thickness of 100 μm were obtained from microfluidic ChipShop GmbH (Jena, Germany). 384-well plastic castings were purchased from Greiner Bio-One GmbH (Frickenhausen, Germany). The following DNA constructs were kindly provided by the indicated persons: HA-RI-α-PARC-GFP (bait-PARC encoding regulatory subunit RI-α of protein kinase A) from Leif Dehmelt (MPI Dortmund, Germany), Gab1-GFP from Fred Schaper (Otto-von-Guericke-University Magdeburg, Germany), CFP-PI3K(p85α) from Shin-Ichiro Takahashi (University of Tokyo, Japan), Shc-RFP from John E Ladbury (University of Leeds, UK), GFP-SOS1 from Giorgio Scita (IFOM Milan, Italy), CFP-HRas from Philippe Bastiaens (MPI Dortmund, Germany), Raf1-CFP from Emilia Galperin (University of Kentucky, USA), MEK1-GFP from Rony Seger (Weizmann Institute of Science, Israel), sfGFP from Peter Pohl (JKU Linz, Austria) and Grb2-YFP from Lawrence E. Samelson (NIH Bethesda, USA).

### Construction of bait-PAR-Grb2

To create arrays of cytosolic Grb2 (bait protein) inside living cells, a HA-PAR-Grb2 construct was generated that transfers the micrometer-scale antibody surface pattern. For this purpose, the regulatory subunit RI-α of the protein kinase A in the previously published HA-RI-α-PARC-GFP (Gandor et al., 2013) was replaced with Grb2 as the following: For seamless DNA insertion, the exponential megapriming PCR (EMP) method was used for all cloning steps (Ulrich et al., 2012). The regulatory subunit RI-α was replaced with Grb2 by amplifying a 800 bp PCR product containing the Grb2 sequence flanked by sequences homologous to the 5’-site and the 3’-site of the HA-RI-α-PARC-GFP vector. The obtained product was purified (QIAquick PCR Purification Kit, Qiagen, Vienna, Austria) and used as a megaprimer in a second PCR run. In a second cloning attempt, the GFP tag was replaced by sfGFP using the identical strategy. Three linker sequences were inserted by round-the-horn PCR. The sequences of interest were divided in two halves and each site was used as a tag on a primer annealing at the respective site where the linker should be inserted. The blunt ends after PCR were ligated by T4 Ligase (Thermo Fisher, Linz, Austria). Linker sequences are depicted in Figure 3D.

### Cell culture and transfection

All cell culture reagents were purchased from Biochrom GmbH (Berlin, Germany). HeLa cells (ATCC) were cultured in RPMI medium supplemented with 10% FBS and 1% penicillin/streptomycin and grown at 37 °C in a humidified incubator with 5% CO2. For transient transfection, cells were sub-cultured the day before and were then transfected with plasmids using the jetOPTIMUS^®^ DNA transfection reagent (Polyplus transfection, Illkirch, France), according to the manufacturer’s instructions.

### Microcontact printing

A PDMS stamp was replica molded by casting PDMS prepolymer mixed in a ratio of 10:1 (component A:B) onto a photolithographically fabricated patterned silicon master. The silicon master (100 mm in diameter) containing a full array of round shaped pillars with a feature size and a depth of 3 μm was obtained from Delta Mask B.V. (Enschede, Netherlands). The PDMS stamp was peeled off the mask and stored at room temperature. The preparation of the micropatterned COP foil was carried out as the following: Briefly, COP foils were washed with ethanol and dH_2_O before hydrophilization by plasma oxidation. Subsequently, hydrophilized COP foils were incubated overnight in GPS/ethanol (1:100, v:v) to form a monolayer of epoxide functional groups on the surface followed by washing with ethanol. For microcontact printing, the large-area PDMS stamp was washed by flushing with ethanol (100%) and distilled water. After drying with nitrogen, the stamp was incubated in 50 mL BSA (or BSA-Cy5) solution (1 mg/mL) for 30 min. This step was followed by washing the stamp again with phosphate-buffered saline (PBS) and distilled water. After drying the stamp with nitrogen, the stamp was placed by its own weight on the clean epoxy-coated COP foil and incubated overnight at 4 °C. The next day, the stamp was carefully stripped from the substrate and the foil was bonded to a 384-well plastic casting using an adhesive tape (3M) and closed with an appropriate lid.

### Live cell micropatterning experiments

For live cell experiments, selected 384-well reaction chambers were incubated with 20 μL/chamber streptavidin solution (50 μg/mL) for 30 min at room temperature. After washing two times with PBS, 20 μL/chamber biotinylated antibody solution (10 μg/mL) was added for 30 min at room temperature. Lastly, the incubation chambers were washed twice with PBS, and cells were seeded at defined cell density for the live cell microscopy analysis. The cells were allowed to attach to the surface for at least 3-4 h prior to imaging to ensure a homogeneous cell membrane/substrate interface, which is a prerequisite for quantitative total internal reflection fluorescence (TIRF) microscopy.

### TIRF microscopy

The detection system was set up on an epi-fluorescence microscope (Nikon Eclipse Ti2). A multi-laser engine (Toptica Photonics, Munich, Germany) was used for selective fluorescence excitation of CFP, GFP, RFP, and Cy5 at 405, 488, 561, and 640 nm, respectively. The samples were illuminated in total internal reflection (TIR) configuration (Nikon Ti-LAPP) using a 60x oil immersion objective (NA = 1.49, APON 60XO TIRF). After appropriate filtering using standard filter sets, the fluorescence was imaged onto a sCMOS camera (Zyla 4.2, Andor, Northern Ireland). The samples were mounted on an x-y-stage (CMR-STG-MHIX2-motorized table, Märzhäuser, Germany), and scanning of the larger areas was supported by a laser-guided automated Perfect Focus System (Nikon PFS).

### TIR-FRAP experiments and calculation of diffusion coefficients

FRAP experiments were carried out on the epi-fluorescence microscope as described above. Single patterns were photo-bleached (Andor FRAPPA) with a high-intensity laser pulse applied for 500 ms. Recovery images were recorded at indicated time intervals. Normalization of data was done by pre-bleach images, and first data analysis was carried out using NIS Elements software package (Nikon). Further data processing was done in Graphpad Prism as described below. Resulting FRAP curves were plotted based on the standard error of the mean and fitted using a bi-exponential equation. Kinetic FRAP parameters were directly obtained from curve fitting.

### Contrast quantitation and statistical analysis

Contrast analysis was performed as described previously (24658383). In short, initial imaging recording was supported by the Nikon NIS Elements software. Images were exported as TIFF frames and fluorescence contrast analysis was performed using the Spotty framework (Borgmann et al.). The fluorescence contrast <c> was calculated as <c> = (F^+^ – F^-^)/(F^+^ – F_bg_), where F^+^ denotes the intensity of the inner pixels of the pattern. F^-^ shows the intensity of the surrounding pixels of the micropattern, and F_bg_ the intensity of the global background. In order to correct for putative differences in bait patterning, results were normalized for bait fluorescence contrast were indicated. Data are expressed as the means ± SE. An unpaired t-test was used to compare two experimental groups. Comparisons of more than two different groups were performed using two-way ANOVA, which was followed by Tukey’s multiple comparisons test in GraphPad Prism software (version 7).

## Acknowledgments

We thank Christian Forsich for technical assistance in plasma activation of COP foils. This research was funded by the province of Upper Austria as part of the FH Upper Austria Center of Excellence for Technological Innovation in Medicine (TIMed CENTER), the Christian Doppler Forschungsgesellschaft (Josef Ressel Center for Phytogenic Drug Research) and the ‘Dissertationsprogramm der Fachhochschule OÖ 2020’ with the financial support of the province of Upper Austria (Austrian Research Promotion Agency (FFG) grant #35409758).

## Author Contributions

P.L. and J.W. designed research; N.O. developed the modified bait-PAR constructs; R.H., U.M., and P.L. performed all experiments; R.H., U.M. and P.L. analyzed data; P.L. prepared the figures and wrote the manuscript. All authors discussed the results and commented on the manuscript.

## Online supplemental material

Figure S1 shows the impact of substrate patterning on bait and prey distribution. Figure S2 shows the investigation of further signalling proteins of the Ras-Raf-MEK pathway.

