## Supplementary Figures 1 and 2 for "Subcellular micropatterning for visual immunoprecipitation reveals differences in cytosolic protein complexes downstream the EGFR"

### Supporting Information

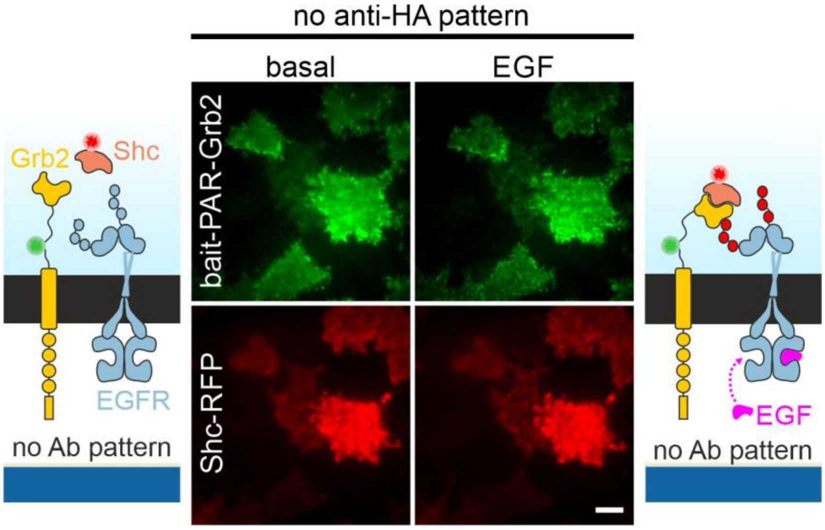


**Figure S1.** Impact of substrate patterning on bait and prey distribution. Cells co-expressing bait-PAR-Grb2 (GFP-labelled) and Shc-RFP were grown on BSA-passivated COP substrates without antibody patterning. Distribution of bait and prey was assessed by TIRF microscopy before and after EGF stimulation (170 nM, 10 min). Scale bar: 15 µm.

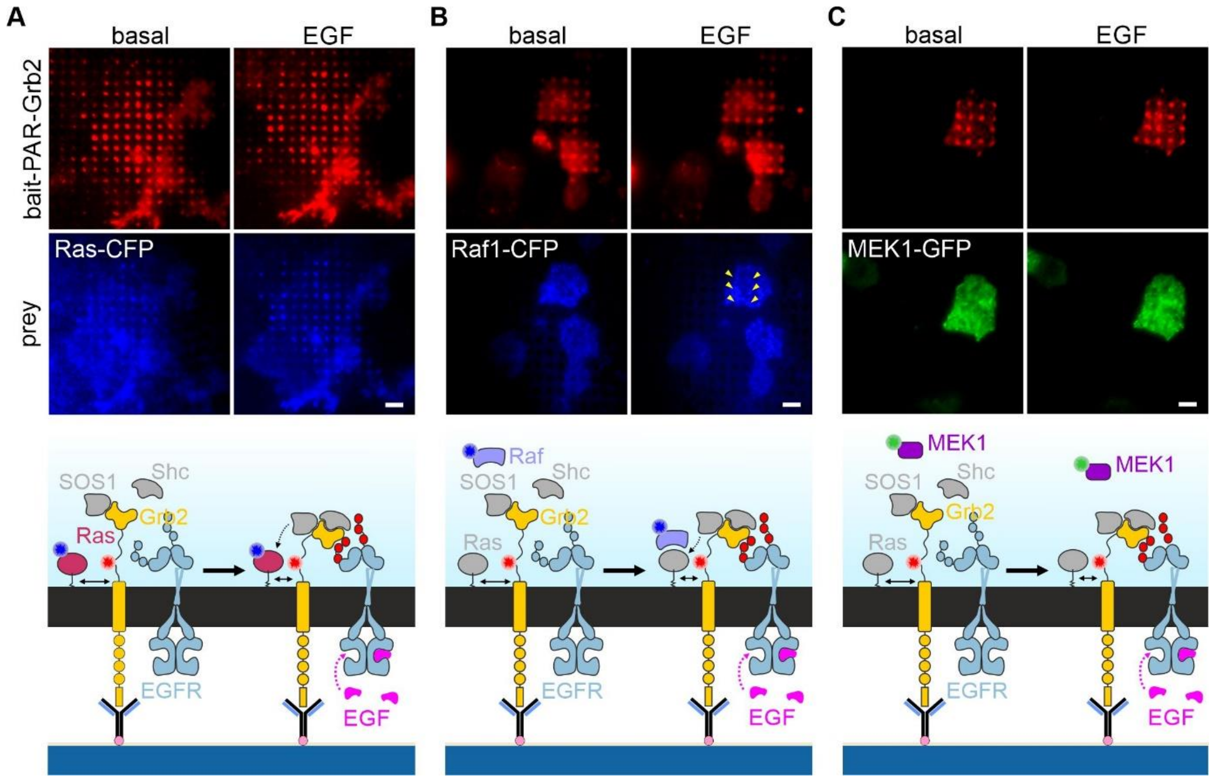


**Figure S2.** Investigating the Ras-Raf-MEK pathway. Representative TIRF microscopy images are shown of cells grown on anti-HA patterned substrates and co-expressing bait-PAR-Grb2 (RFP labelled) and (A) HRas-CFP, (B) Raf1-CFP, and (C) MEK1-GFP. Scale bar: 15 µm. Schematic presentations illustrate Ras-Raf-MEK downstream signalling pathway and formed protein complexes.
